# The ENIGMA MEG Pipeline: Automated cortically localized spectral analysis of multi-site resting state MEG datasets

**DOI:** 10.64898/2026.06.03.729876

**Authors:** Allison C Nugent, Anna M Namyst, Frederick W Carver, Paul M Thompson, Jeffrey D Stout

**Affiliations:** Magnetoencephalography Core Facility, National Institute of Mental Health, National Institutes of Health, Bethesda, MD, United States; Imaging Genetics Center, Mark C Mary Stevens Institute for Neuroimaging and Informatics, Keck School of Medicine, University of Southern California, Los Angeles, CA, United States

**Keywords:** Magnetoencephalography, resting state, power spectral density, FAIR principles

## Abstract

**Background:** Magnetoencephalography (MEG) is a unique technique in human neuroimaging combining high temporal resolution (millisecond or faster) with moderate spatial resolution (several millimeter). While many software packages for MEG data analysis exist, there is no pipeline developed for the specific purpose of enabling the automated analysis of very large, multi-site datasets.

**Results:** The ENIGMA consortium was developed to enable large scale collaborations in the fields of neuroimaging and genetics. To facilitate ENIGMA MEG working group data analysis, we developed the ENIGMA MEG pipeline. The first ENIGMA MEG working group project involves spectral analysis of resting state MEG data, thus our current pipeline is designed to carry out that task. The goals of the ENIGMA MEG pipeline include ease of use, automated processing wherever possible, detailed logging and quality assurance (QA) features, the use of the brain imaging data structure (BIDS) format, anonymized output, and consistent processing across vendors. The pipeline is built using the MNE-Python framework and incorporates a re-trained version of the MEGnet deep neural network algorithm for automated artifact detection. QA tools are designed to enable high throughput evaluation of a large number of subject datasets. All software is open source and available on GitHub (https://github.com/nih-megcore/enigma_MEG). We used our pipeline to process data from three publicly available MEG cohorts, demonstrating its functionality and compatibility with large-scale processing.

**Conclusions:** While the current ENIGMA pipeline is limited to resting state data and spectral analysis for the current working group project, the software is highly modularized, allowing straightforward extension to other analysis questions. Further development of the tool to enable connectivity and task-based MEG analysis are planned. The ENIGMA MEG pipeline represents an important first step to augment the existing arsenal of analysis tools, enabling multi-site, high throughput data analysis.

## Introduction

Single site data collection and analysis is typical in most scientific studies, however, this often results in a lower number of recruited subjects and a bias towards site specific effects [1–3]. Several publications have demonstrated a dearth of reproducibility in science overall and neuroimaging studies specifically [4–6] - often referred to as a reproducibility crisis. While inadequate power is a consistent problem across the literature, power analyses are rarely performed [7–9]. Some studies, such as brain-wide association studies (BWAS), require subject numbers in the thousands to prevent falsely inflated correlations and inferences [10]. Such a large sample size is rarely achieved in the literature and is almost impossible for a single site study. In response to these issues, the fMRI community has produced many large, multi-site data collections, including the Adolescent Brain Cognitive Development study (ABCD; [11]), with over 10,000 MRI scans; the Alzheimer’s Disease Neuroimaging Initiative (ADNI; [12]), with over 17,000 structural MRIs; and the UK BioBank (the 2020 release included over 35,000 MRI scan participants [13]). While there are some large MEG cohorts available, the largest being the Cambridge Centre for Ageing and Neuroscience (Cam-CAN) study [14], with over 600 participants, none reaches the magnitude of the largest available MRI data collections. In addition, multi-site MEG studies are rare, with several notable exceptions. The Open MEG Archive (OMEGA), including 644 participants (444 healthy controls and 200 patients) [15];the MEG-UK data collection includes over 400 datasets from healthy controls across eight sites [16]; the BioFIND data collection includes 324 datasets collected at two sites in patients with mild cognitive impairment and healthy controls [17]; and the Dev-CoG study includes longitudinal data from over 200 subjects at two sites [18].

Many factors have led to the lack of multi-site MEG studies. MEG is a less common neuroimaging modality with fewer research sites engaged in data collection compared to brain MRI. In addition, MEG data analysis across multiple sites has been further limited in part because of differences in sensor types, vendor preprocessing steps, data formatting, and data layout. One particular area of difference between vendors is in how environmental noise is mitigated. MEGIN/Elekta scanners use a software signal processing method incorporating site-specific calibration files [19]. While effective, the procedure dramatically reduces the rank of the underlying raw data. In contrast, the other manufacturers use arrays of reference sensors placed at some distance from the primary sensors, although each manufacturer differs in how the signals from the reference sensors are used to correct for external noise [20]. The lack of standardized processing pipelines across vendors has also limited progress in the MEG field. The ENIGMA Consortium has emerged as a network of researchers working on large scale data analysis projects [21]. As part of this initiative, the ENIGMA MEG Working Group was created, in an effort to overcome these challenges and bring together as many MEG sites as possible in a collaborative effort to assemble and analyze large, multi-site MEG data collections. The end goal is to increase statistical power and produce reproducible, generalizable, and clinically relevant results.

The analysis of very large numbers of datasets does come with its own set of unique challenges. In particular, computing outputs and performing quality assurance (QA) on the results requires a large investment in time, so an automated pipeline is crucial to routine use of a given pipeline and the reduction of variance between user outputs. To enable the systematic analysis of large cohorts, the fMRI neuroimaging community has built many automated data analysis pipelines with robust QA measures to reduce the data analysis burden; four notable examples are fMRIPrep [22], HALFpipe [23], DCAN [24], AFNI_proc [25], and the Human Connectome Project (HCP) pipeline [26]. While most of these tools focus on single subject processing, group level analysis of very large fMRI datasets has been integrated in the Image-based-meta- and mega-analysis (IBMMA) tool [27]. In contrast, there have been a relatively limited number of automated pipelines available for MEG analysis. Recently, the FLUX pipeline was released, with the goal of standardization and training, although data analysis using the pipeline still requires substantial manual intervention [28], and was thus inappropriate for the analysis of very large datasets. Two other options that allow for more automated processing are the OSL pipeline [29] and MNE-BIDS-Pipeline [30]. There are several reasons why these packages did not fully meet our needs. The MNE-BIDS-Pipeline does not support beamforming source modeling, which we felt was crucial for artifact rejection. Artifact removal using ICA requires manual identification of artifactual components, while comprehensive output reports are generated, there is no facility for high throughput screening of large numbers of reports or QA images. While the OSL pipeline will perform automatic artifact component removal, it requires the presence of EOG and ECG channels in the data, which are not collected at all (or even most) sites. The motivation for the ENIGMA MEG pipeline was to create a self-contained, fully-automated processing stream for high-throughput data analysis, capable of uniform processing of data from all MEG imaging platforms. Because the initial ENIGMA MEG working group project investigates spectral characteristics in resting state data, the initial release of the ENIGMA MEG pipeline focuses on this goal. However, because future analyses are planned, we sought to design the software using a modularized coding approach that would allow for future integration of other analysis methods.

One issue with processing multisite datasets, particularly if not all datasets will be processed at a single location, is uniformity of data organization. The gold standard for data organization in the neuroimaging community is the brain imaging data structure (BIDS) standard [31]. The BIDS standard requires a specific file structure, naming convention, and additional json format sidecar text files that contain descriptive elements. The BIDS standard has been extended to MEG, incorporating all metadata relevant to data analysis [32]. Importantly, in the BIDS specification where MRI data is shared along with the MEG data, anatomical landmarks indicating the position of the MEG fiducials in the space of the anatomical MRI must be included in the sidecar files of the MRI data and/or the sidecar file describing the MEG coordinate system (*_coordsystem.json). If this information is not shared, then the MRI and MEG data cannot be properly coregistered and source localization is not possible. The MNE-BIDS module was developed to facilitate conversion of MEG and associated MRI data to BIDS format and is used within the ENIGMA MEG pipeline to manage file access and output [33]. The primary analysis functions are taken primarily from the MNE Python package.

Due to the large number of sites and individual datasets involved, and the desire to be maximally inclusive to a variety of sites, we designed the ENIGMA MEG pipeline to be fast and easy to use for researchers with varying levels of expertise in data analysis. These objectives resulted in an easy-to-use conversion to BIDS, automated data analysis, understandable logging, and GUI based quality assurance. Automated artifact detection is implemented through integration of the deep learning MEGnet algorithm [34], retrained on data from the major MEG manufacturers. The project level, panel-based QA process is intuitive, because it relies on our visual system’s natural outlier detection process. Ultimately this results in a simplified end-to-end solution for MEG processing that removes the time burden on the researcher. As currently written, the ENIGMA MEG pipeline is not a general-purpose tool; it is specifically intended for spectral processing of resting state MEG data. While it was developed in the context of the ENIGMA MEG Working Group, it can be used to process any MEG resting state data. It is written in a modular format, however, allowing for extension to other analysis types. We expect that most, or all future projects of the ENIGMA MEG working group will use extensions to the original ENIGMA MEG pipeline. This manuscript provides an overview of the current software, shares our design philosophy, and hopefully demonstrates a framework that can be developed in the future to address other analysis tasks.

## Methods

### Basic Architecture

The ENIGMA MEG pipeline is built in Python, largely using functions from the MNE-Python package, and is available on GitHub [35]. Full installation instructions are given in the documentation. We recommend that dependencies (MNE Python) are installed through conda or mamba, then the ENIGMA MEG pipeline code is installed via pip. The primary processing function, process_meg.py (Figure 1), is built around the process class, with individual methods to carry out each step in the analysis. Thus, the pipeline is easily extensible to different analysis goals. Additionally, as new methods are developed, additional processing methods can be easily incorporated into the pipeline using the current architecture. We will discuss potential additional processing models that could be added to increase functionality for resting state data in detail later in the Methods section. Following processing, there are additional routines to generate images for QA review. As currently written, there are two requirements to use the pipeline. First, the raw data must include a resting state MEG scan and a structural MRI scan, organized in BIDS format. Given that the pipeline is designed to operate on multi-site data, some standardization of input formatting is crucial. While not all metadata options in the BIDS structure need to be supplied, the fields giving the coordinates of the MEG head localization coils in the anatomical space of the MRI are required. A separate package, enigma_anonymization_lite [36], available on GitHub, was constructed to assist in forming a BIDS directory for MEG and MRI data co-registered using MNE-Python. This routine is a lightweight wrapper around the MNE BIDS module [32], allowing for .csv file inputs and QA outputs, that additionally performs anonymization of the MEG data and MRI scan to facilitate data sharing. This function allows MNE-Python and Brainstorm users to easily convert their data into a minimal BIDS format that allows processing with the ENIGMA MEG Pipeline. The second requirement is that the MRI must be processed with the FreeSurfer recon-all pipeline through the cortical parcellation steps [37, 38]. While this requires additional time from the user, FreeSurfer is a fully automated procedure, and the ENIGMA MEG Pipeline produces diagnostic information on the quality of FreeSurfer segmentation in the logfile, and the QA tools allow for rapid QA of segmentation. An additional empty room MEG recording is optional, but can be included if available. Further details of the pipeline are given in the sections below. The package also contains helper functions designed to parse an existing BIDS directory for resting state datasets and to create a text file which can be used for batch processing (for example, using the slurm sbatch --array function) on a cluster.

**Figure 1:**
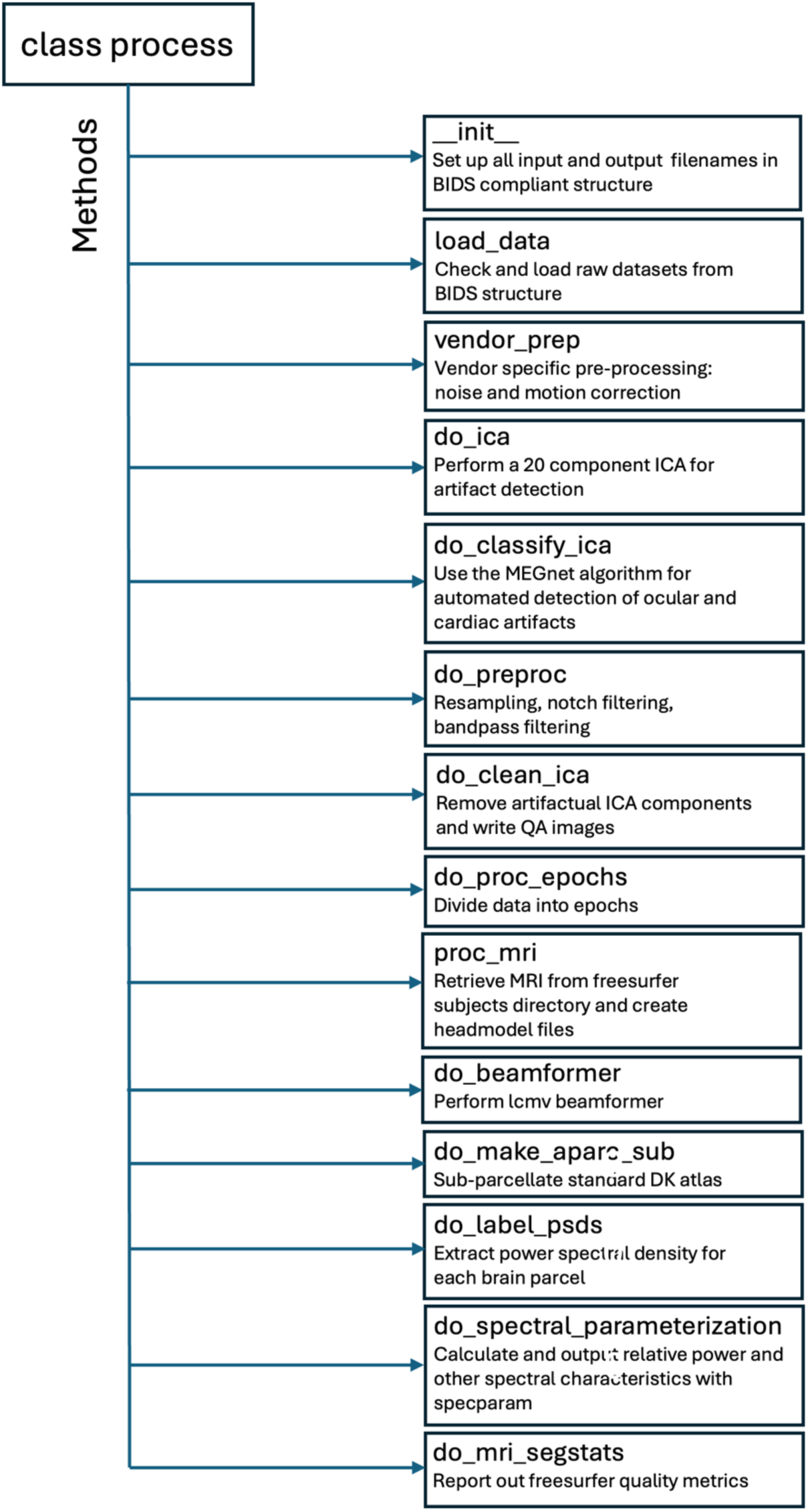
Overview of the structure of the process class, the heart of the ENIGMA MEG pipeline. Listed are currently implemented methods for each stage of processing. This modular design allows for easy extension to additional analytic tasks.

### Initial Preprocessing

The first data processing segment of the ENIGMA pipeline performs initial preprocessing in a vendor specific manner. Our goal was to perform vendor specific pre-processing only, such that all datasets regardless of platform could be processed using the same analysis steps and parameters. For CTF datasets, 3^rd^ order gradient compensation is applied for noise correction, then both the resting state and empty room data are assessed for bad channels using the MNE-Python find_bad_channels_maxwell function, plus an additional algorithm based on the standard deviation to identify flat channels. MEGIN/Elekta data is first checked to see if Maxfilter has already been performed. If not, bad channels are identified, then the MNE-Python implementation, maxwell_filter() is performed, with movement correction if continuous head position data exists [19]. For other vendors, only the assessment of bad channels is performed.

### ICA artifact detection

The next step in processing is artifact correction, using independent components analysis (ICA). First, a twenty component ICA is performed using MNE-Python (after downsampling to 250Hz, powerline filtering, and 1-100Hz bandpass filtering). These parameters were chosen to align with the preprocessing of the data used in the original MEGnet investigation. For each component, topographical maps are interpolated to a circular image. The twenty topographical images and source time series are then classified using the MEGnet algorithm [34], which uses a convolutional neural network model for automated classification of both spatial and temporal components. The MEGnet model architecture was optimized using a 20 component ICA, the use of a different rank could potentially degrade performance [34]. The initial publication states that other component numbers were tested, and a 20 component ICA was empirically judged to produce ICA components most easily identified by human raters. The original MEGnet algorithm was trained using 232 MEG recordings collected from 171 subjects acquired on either a CTF system or a 4D system. To enable (and validate) artifact detection from all four major vendors, we used the original model architecture but reset the layer weights to a uniform distribution between −1 and 1 and performed training from scratch using a new data collection described below. The training was performed on an Nvidia RTX 6000 with 48 Gb of GPU RAM, with input from the original authors. The number of epochs was 20, the batch size was 300, and we used the Adam optimizer with a learning rate of 1e-3. The imbalanced class weights were computed from the data using sklearn and entered into the model fit. The data collection used for training consisted of 896 MEG recordings from 489 subjects scanned on 4D/BTi, KIT, MEGIN/Elekta, or CTF platforms. The data was processed using ICA, and components were manually classified by one of three raters. Two raters had extensive experience in MEG and manual identification of ICA components (co-authors ACN and AMN), and a third worked under the supervision of the other two. In most cases, eyeblink, saccade, and cardiac artifacts were readily identifiable based upon canonical spatial topographies and time series artifacts. Any edge cases were flagged and discussed by at least two raters and a consensus identification was made. Because a consensus-based format was used, where raters were allowed to consult eachother to achieve the most accurate rating, calculation of interrater reliability was not appropriate. However, we additionally had two raters identify components in an additional set of 50 recordings from participants with epilepsy. Although the same consensus rating method was still used, we had both raters rate all subjects to verify that the consensus method was valid. For components rated as high confidence by both raters (i.e. not flagged for adjudication), Cohen’s kappa was 0.96 (raw agreement 99.3%), indicating excellent agreement.

Both resting state and working memory recordings from the human connectome project (HCP [39]) were used, along with 445 recordings in one of several tasks (including resting state) from the NIMH Healthy Volunteer cohort [40]. Additionally, a subset of 245 resting state recordings from the larger CAMCAN data repository [14] were used, as well as 45 resting state recordings acquired on the KIT scanners at NYU in Abu Dhabi and New York (personal communication, Flick, G., Pylkkanen, L., and Marantz, A.). All 896 MEG recordings were used for the MEGnet model training. A holdout set of 20% (N=184) of the data was selected from the total using the MultilabelStratifiedKFold function from the iterative-stratification package [41] to align the test/train and holdout distribution on scanner, acquisition site, age, gender, and task type. A breakdown of the training/test data across scanners is given in Table 1. The test/train sample was broken into a 1:6 ratio of testing (N=103) and training (N=609), and the model weights derived from training were then used to assess the holdout sample. The accuracy of the algorithm in the holdout sample is shown in Table 2. Similar to the original MEGnet, our model was 98.78% accurate at classifying independent components. The most common error was to classify non-artifactual components as cardiac associated; of 3,230 non-artifactual components, 28 were classified as cardiac artifacts (0.87%). In our supplementary materials (Tables 1-7), we additionally give confusion matrices stratified by site and by task (eyes open rest, eyes closed rest, or other task). While some differences in individual artifact class false positives were noted across sites, all sites performed with accuracy > 98%. Because subjects from two sites (NIH, HCP) had multiple scans, these were split across test and train sets for some subjects. Notably, however, sites with only one recording per subject (CAMCAN, NYU) showed equally high accuracy (Accuracy for the CAMCAN set was highest at 99.07%). MEGnet compares favorably to other similar published methods for automated ICA component removal, although none of those tools are available publicly [42–44]. The retrained model and associated implementation software is available on GitHub [45], and is installed automatically as part of the ENIGMA pipeline installation. Additional testing in a sample of participants with epilepsy revealed that the model trained on healthy volunteers tended to over-identify artifactual components in the epileptic sample; thus, we recommend additional training of the model would be required for that patient group. We would expect that the model is generalizable to healthy volunteers or patients with other disorders that would not be expected to produce gross abnormalities in the MEG signal. We would not recommend using our trained model on other scanner configurations without additional testing, or model retraining if necessary.

**Table 1:** Datasets used in MEGNET re-training, along with number of subjects, number of datasets, and breakdown between test/train and holdout datasets.

| Dataset | Vendor | Total Subjects | Total Datasets | Training Datasets | Holdout Datasets |
| --- | --- | --- | --- | --- | --- |
| HCP | 4D/BTi | 88 | 161 | 130 | 31 |
| NYU | KIT | 45 | 45 | 35 | 10 |
| CAMCAN | MEGIN/Elekta | 245 | 245 | 186 | 59 |
| NIH | CTF | 111 | 445 | 361 | 84 |
| Totals |  | 489 | 893 | 712 | 184 |

**Table 2:** Distribution of components identified in the data from manual classification (actual class) as well as the distribution of predicted class from the re-trained MEGnet algorithm. We also show Sensitivity, False Negative Rate, Positive Predictive Value, and False Discovery Rate. Overall accuracy and error rate appear in the lower right.

|  | Predicted Class |  |  |  |  |  |  |
| --- | --- | --- | --- | --- | --- | --- | --- |
|  |  | Non-Artifact | Blink | Cardiac | Saccade | Sensitivity | False Negative Rate |
| Actual Class | Non-Artifact | 3185 | 9 | 28 | 8 | 98.61% | 1.39% |
|  | Blink | 0 | 179 | 0 | 0 | 100.00% | 0.00% |
|  | Cardiac | 0 | 0 | 195 | 0 | 100.00% | 0.00% |
|  | Saccade | 0 | 0 | 0 | 76 | 100.00% | 0.00% |
|  | Positive Predictive Value | 100.00% | 95.21% | 87.44% | 90.48% | Accuracy | 98.78% |
|  | False Discovery Rate | 0.00% | 4.79% | 12.56% | 9.52% | Error Rate | 1.22% |

### Additional preprocessing

Following the ICA procedure, additional preprocessing of the data takes place. First, data is resampled to 300 Hz, and notch filtering is performed at the mains frequency (50 or 60 Hz, depending on the origin of the data). Data is additionally bandpass filtered between 1 and 45 Hz using the default filter in MNE-Python: a zero-phase overlap-add FIR filter using a Hamming window. The underlying filter is designed using scipy.signal.firwin(). Further information on default methods for filtering are available on the MNE-Python website. Note that the 45 Hz cutoff was chosen specifically in response to the intention of using the pipeline in combined cohorts where some of the datasets would have a 50Hz powerline frequency artifact, while other datasets would have a 60Hz artifact. While notch filtering at the mains frequency is standard practice, we were concerned about injecting potential bias into any subsequent analysis of the 45-65Hz bandpass due to the differing frequency contributions from different sites, which motivated our cutoff at 45Hz. For users who intend to analyze data of only a single mains frequency, modification of the code to allow alternative implementations is relatively straightforward in our open-source codebase. Following filtering, the ICA components previously identified as artifactual are projected out of the data. Next, the data is divided into four second epochs. Epoch lengths in the range of 2-4 seconds are common in literature exploring spectral power [46, 47]. Epochs are rejected if the peak-to-peak signal amplitude of a magnetometer channel exceeds 5000 fT or a gradiometer channel exceeds 5000e fT/m. Note that a liberal upper threshold was chosen because we expected that the ICA cleaning and use of an LCMV beamformer would provide the bulk of the artifact removal, and our intention was to retain the maximum amount of data. Nevertheless, these thresholds will remove large muscle artifacts and sensor jumps. Epochs are also excluded if the peak-to-peak signal amplitude of a magnetometer channel is below 10 fT or a gradiometer channel is below 10 fT/m. Given that the noise floor of a SQUID sensor is generally around 5 fT, any signal not exceeding a peak-to-peak signal amplitude of 10 fT is likely to be unreliable; in our experience even a low-signal pediatric subject will easily exceed this threshold provided the participant is correctly positioned in the helmet. The final number of epochs and total time is recorded in the logfile.

### MRI preprocessing

Before running the ENIGMA processing pipeline, the anatomical MRI must be preprocessed using FreeSurfer software [37]. The extra MRI processing steps occurring as part of the pipeline create the head model required for source localization. First, the MNE-Python routine make_watershed_bem uses the FreeSurfer watershed algorithm to create boundary element model (BEM) brain, inner skull, outer skull, and outer skin surfaces. These surfaces are in turn used to create the surface-based source model and forward solution using the coregistration information embedded in the BIDS metadata structures. We used a 448 parcel sub-parcellation of the standard FreeSurfer Desikan-Killianay (DK) atlas that was designed specifically for MEG studies [48]. While this parcellation has been used for MEG functional connectivity, the number of parcels is larger than the sensor count in any commercial MEG system; thus, parcels are necessarily non-independent. This is exacerbated by algorithms such as MaxFilter that necessarily dramatically reduce rank. Since this parcellation is a sub-parcellation of the DK atlas, and the original DK parcel names are retained, the 68 parcel segmentation can be recovered from the output files.

### Source Localization

After MRI processing is completed, source localization is performed using a linearly constrained maximum variance (LCMV) beamformer, calculated on every vertex on the surface. An LCMV beamformer was chosen specifically because of its ability to effectively suppress artifacts [49], crucial in multi-site studies where data quality may be non-uniform across sites. The choice of an LCMV beamformer over a DICS (dynamic imaging of coherent sources) beamformer was motivated by several reasons. First, more recent studies with similar goal to the ENIGMA MEG project utilized LCMV beamformers, including Rempe, et al. [47]and Doval, et al. [50] (although one notable exception, Stier, et al., did utilize a DICS beamformer [51]). In addition, a recent study investigating power maps using the FOOOF/specparam algorithm also utilized an LCMV beamformer [46]. Because we wanted to output the entire spectrum, we felt that it was important to use the same spatial filter over the entire spectrum, rather than compose a spectrum from narrow-bandwidth DICS estimates. Finally, we felt that an LCMV beamformer offered increased flexibility, considering that we want to allow for additional methods in the codebase for additional hypotheses. We did originally test a broadband DICS beamformer to allow for frequency domain estimation of the spectrum, but found it to be significantly less stable, sometimes failing to produce reasonable source maps. Nevertheless, this option remains in the code.

First, a broadband covariance matrix is calculated for both the resting state and empty room recordings. In cases where an empty room recording is not available, an ad hoc covariance matrix is used for normalization. Regularization of 5% is applied (this is the default value in MNE-python [52]). Small amounts of regularization dramatically increase power, signal to noise, and accuracy of the reconstructed signal [53]. While adaptive regularization is possible, we were concerned that this could potentially introduce bias, if sites differed substantially both in mean regularization applied and covariates of interest (such as age). Orientation of each source point is determined by the direction that maximizes power, and beamformer weights are normalized using unit-noise-gain. Beamforming parameters were previously tested for all four vendors to ensure uniformity of behavior across platforms [54]. Following the calculation of the beamformer weights, a power spectral density (PSD) is extracted for each parcel in the sub-parcellation by using the 10% trimmed mean of a 15 component principal components analysis (PCA) on the parcel vertices time series [46].

### Output

Power in the five canonical bands is calculated from the PSD in each parcel, and normalized to relative power by dividing by the total power in the 1-45Hz range. In addition, the FOOOF/specparam algorithm [55] is used to extract the peak frequency within the alpha band, as well as the offset and exponent describing the aperiodic component of the PSD. Note that the FOOOF algorithm is performed on the unnormalized spectrum, and the aperiodic offset will thus reflect any global scaling (in addition to parcel specific scaling), and thus may not be appropriate to analyze according to subject characteristics. These parameters are then output for each parcel in a comma separated values (.csv) format file. Additionally, a separate .csv file is created containing the raw unnormalized spectra in every parcel. This ensures maximum flexibility, enabling the user to perform any desired normalization methods, spectral modeling, or alternative calculations of spectral power. Referring to Figure 1, as currently configured, the main pipeline performs extraction of the spectrum and parametrization following beamforming and parcellation (do_beamformer and do_make_aparc_sub). Additional methods could be added here; for example, a “do_connectivity” method, to extract a connectivity matrix between parcels, using amplitude or phase-based methods. Alternatively, a “do_nonlinear” method could potentially be added to calculate complexity or nonlinear dynamics metrics such as Lempel-Ziv complexity, Hurst exponent, or detrended fluctuation analysis. Another option would be a “do_phaseamplitude” method to additionally perform phase-amplitude coupling analysis. These different methods could be invoked at runtime using command line flags, with appropriate checks to see what intermediate files are already available to avoid duplicate runs of prior steps as new methods are added.

### Logging and Quality Assurance

The start and stop of every major subroutine in the processing pipeline is recorded in a logfile. In addition, the logfile retains information that can be used for quality assurance, including the number of ICA components removed, the number of epochs removed (and the total time remaining), and the quality of the FreeSurfer segmentation. An example logfile for a dataset that completed processing is shown in Figure 2. Additional routines produce QA images for review. These include images of the head surface and sensor array to assess coregistration, the boundary element model surfaces used for the calculation of the forward model, the source model, the reconstructed FreeSurfer cortical surface, the average PSD across all parcels and epochs, and an image of relative alpha power plotted in all parcels across the brain. A summary of QA images appears in Figure 3. In addition, in Figure 4 we show the QA GUI interface for rapid identification of data quality issues.

**Figure 2:**
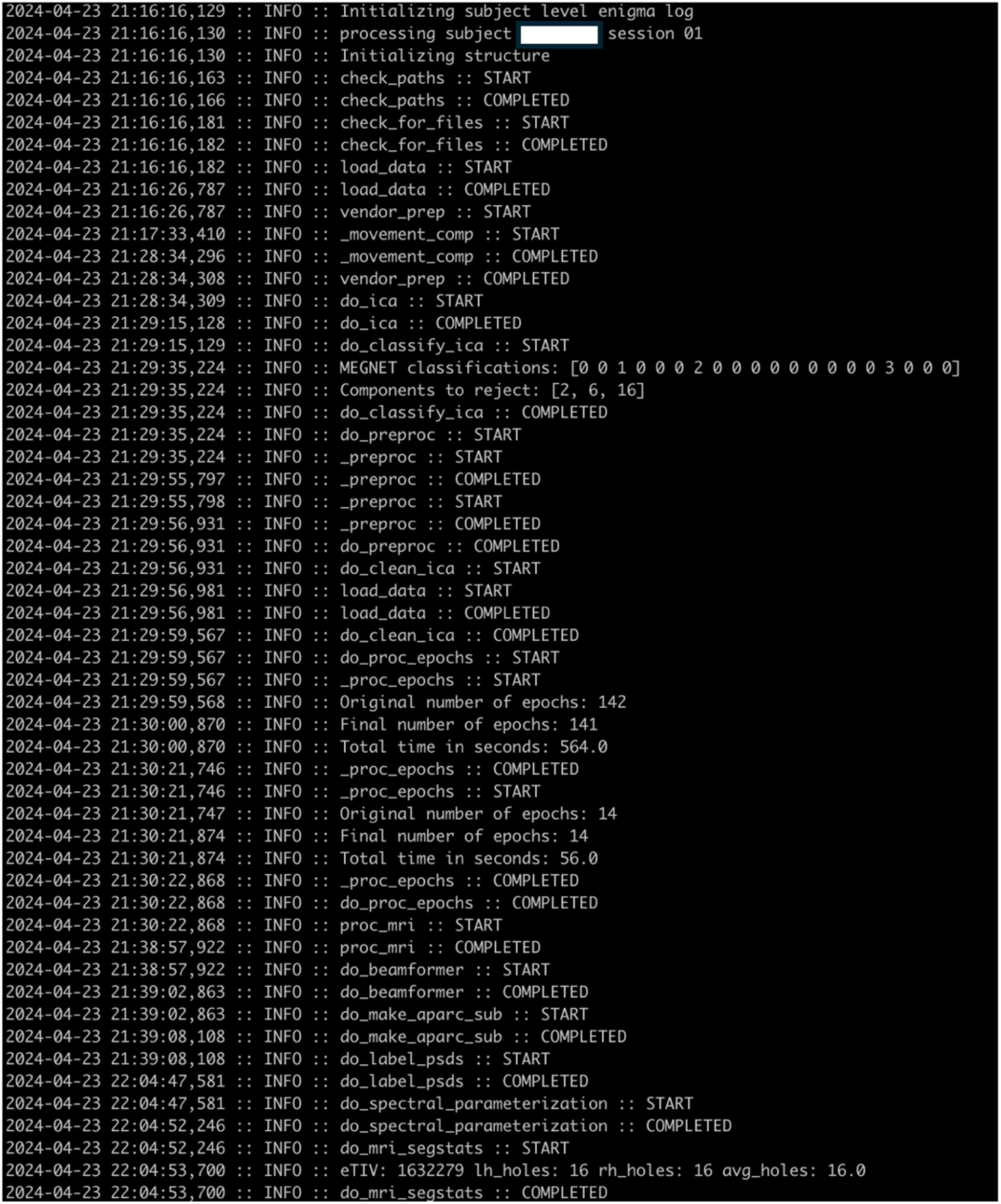
Example log file for a completed dataset. Start and completion of all major routines is indicated for efficient identification of failure point. Quality assurance meta data is available in the log file and can be easily parsed to filter datasets or compile statistics.

**Figure 3:**
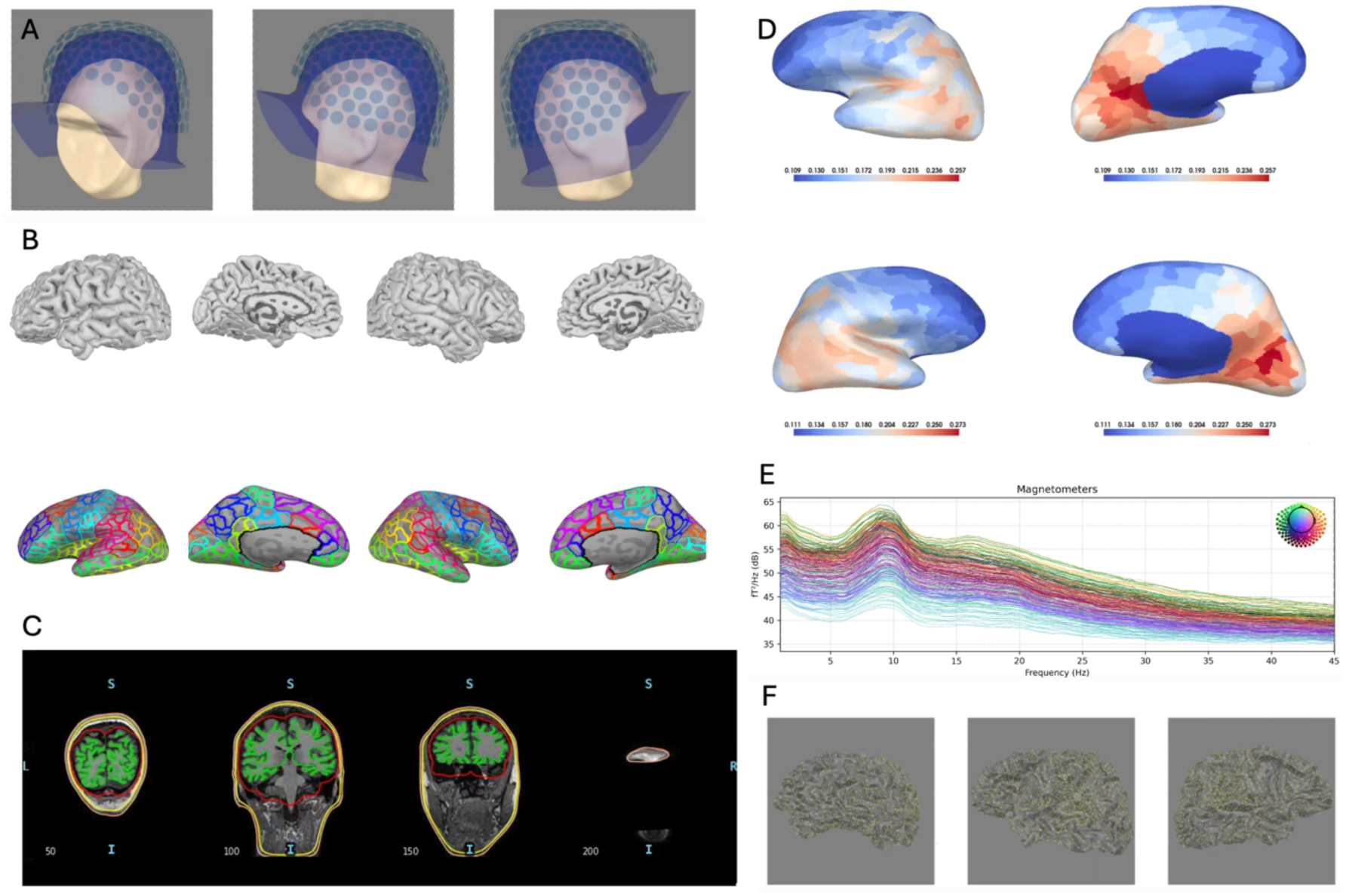
Examples of QA images produced by the ENIGMA pipeline. A) Coregistration of MRI and MEG, B) Cortical extraction and parcellation, C) Boundary element model (BEM) surfaces used for head model creation, D) Relative alpha power plotted across parcels, E) Full power spectrum from all sensors, and F) Source space.

**Figure 4:**
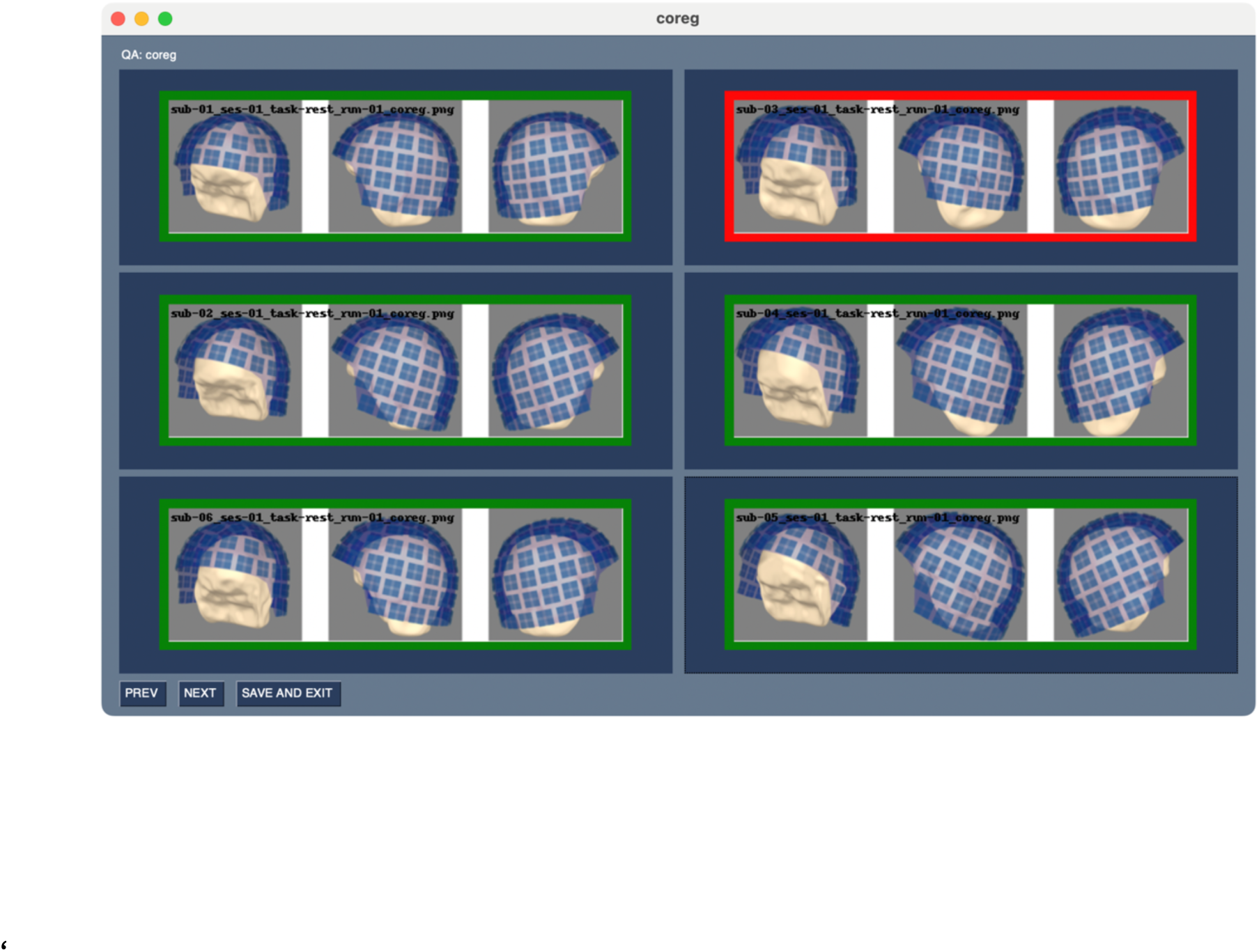
Example of QA viewer to assess coregistration. Images to be retained or omitted can be indicated with a mouse click, using green (include) and red (exclude) color coding. Previous and next buttons are used to display a new panel of images. The number of images displayed is user adjustable to accommodate differing monitor sizes (Image flagged in red for illustration purposes only).

### Demonstration of Functionality in Large Cohorts

The ENIGMA project and analysis was reviewed by the NIH Combined IRB and found to be Non-Human Subjects Research. To provide an example of the final output of the pipeline, we briefly present results from three publicly available cohorts: the Human Connectome Project (4D scanner [39]), the MOUS dataset (CTF scanner [56]) and the Rempe/BoysTown dataset (MEGIN/Elekta scanner [47]). The HCP dataset requires signing a data use agreement to access the data. All three cohorts required additional processing to either convert the data to the BIDS format, or to add metadata required by the ENIGMA pipeline; this additional software is available in the extras/ subfolder of the GitHub repository. In addition, because the HCP datafiles had the locations of the MEG fiducial coils scrubbed, a separate hcp_process_meg.py script was required (also available in the extras/ subfolder). This version of the pipeline utilizes transform files rather than the locations of the MEG fiducial coils in the anatomical space of the MRI. We provide some comparisons between the three cohorts but note that no data harmonization is performed as part of this demonstration (although harmonization will be discussed in detail in the discussion). Because spectral characteristics are strongly dependent upon age, and our cohorts differed in their age distributions, we present comparisons of the full datasets as well as a subset of the Rempe cohort truncated to contain only ages 18-35. We show distributions of spectral power across all subjects and parcels in each frequency band and for the aperiodic exponent. In addition to distributions, we calculate the mean value for each parcel within the datasets and perform Pearson correlations as a measure of spatial correspondence. Finally, we use additional recordings collected in the same session in the HCP cohort to demonstrate test/re-test reliability.

## Results

The HCP cohort included 88 subjects, the MOUS cohort included 199 subjects, and the Rempe/BoysTown cohort included 388 subjects. Of the HCP datasets, one had only a partial MEG recording, and one failed FreeSurfer processing, leaving N=86 datasets that completed ENIGMA pipeline processing. One of the MOUS datasets failed in FreeSurfer processing, and one of the Rempe datasets failed due to data quality issues, leaving N=198 and N=387 completed datasets, respectively. Using a 6 core partition on a single node on the NIH Biowulf cluster (AMD EYPC 9454 48-Core processor), a single MEGIN/Elekta dataset from the Rempe cohort required 8 minutes 32 seconds to complete, while a single CTF dataset from the MOUS cohort required 13 minutes 57 seconds to complete. Note that the Rempe datasets were shared after MaxFilter was performed, Raw MEGIN/Elekta datasets require additional time. A summary of the age and sex makeup of these cohorts, and some associated quality assurance measures are given in Table 3.

**Table 3:** Basic statistics for the three exemplar datasets. Age and gender breakdown are given. Surface holes is a measure of quality of the freesurfer segmentation; datasets with a value over 110 are eliminated from further analysis. Additionally, mean number of 2s epochs is given, both in the original dataset and after epochs were dropped for data quality issues.

|  | Mean Age<br>(range) | F/M sex | Surface holes<br>average<br>(range) | Mean Number of Epochs (range) |  |
| --- | --- | --- | --- | --- | --- |
|  |  |  |  | Original | Post-Processing |
| HCP | 28.4 (22-35) | 47/39 | 48.3 (25-82) | 93.2 (91-115) | 89.5 (56-115) |
| Rempe | 25.4 (6-85) | 204/183 | 23.3 (3-130) | 88.3 (74-135) | 81.0 (1-134) |
| MOUS | 22 (18-32) | 101/97 | 32.5 (5-135) | 80.5 (77-112) | 77.8 (51-110) |

Previous studies have used the reported “average holes” metric, which is related to the Euler characteristic, to serve as a measure of the quality of a FreeSurfer segmentation [57]. Based upon Rosen, et al. [58], we used a threshold of 110 or fewer average holes. In addition, we only retained datasets with at least 10 epochs of usable data, such that the final counts were N=86 (HCP), N=197 (MOUS), and N=382 (Rempe). Note that the datasets with the most epochs removed tended to come from the youngest participants (which were part of the Rempe cohort), which is to be expected due to increased propensity for head motion in this cohort. Note that these values are not hard-coded into the ENIGMA-MEG Pipeline, data is produced regardless of epoch count, or Euler characteristic. These values are reported in the log files so that the user can determine their own threshold for data retention.

Figure 5 shows the mean raw power spectral density over all subjects, with each trace corresponding to a separate parcel. Note that these spectra are not normalized in any way and represent the raw output spectra of the pipeline. The output spectra span the interval 1-45Hz, in 0.25 Hz steps. In Figure 6, we show distributions of data over all parcels and all subjects for each cohort. For the Rempe cohort, we use a subset of the data (N=58) in the age range 18-35, to overlap with the other two cohorts, given the strong dependence of spectral characteristics on age. We then average each cohort over all subjects within each parcel to obtain mean power maps. In Figure 7, we show the relative power in each of the canonical bands, along with the aperiodic exponent, plotted in every parcel for the three cohorts. Note that these are plotted on independent min-max scales for each cohort given the lack of harmonization and adjustment for differences in condition; while the HCP and MOUS recordings were acquired in an eyes-open condition, the Rempe recordings were acquired with eyes closed. For instance, Rempe datasets showed higher alpha power overall compared to the HCP and MOUS datasets, consistent with the well documented increase in alpha power in the eyes closed condition. Alpha power images recapitulate the familiar anterior-posterior gradient in alpha power previously reported [46].

**Figure 5:**
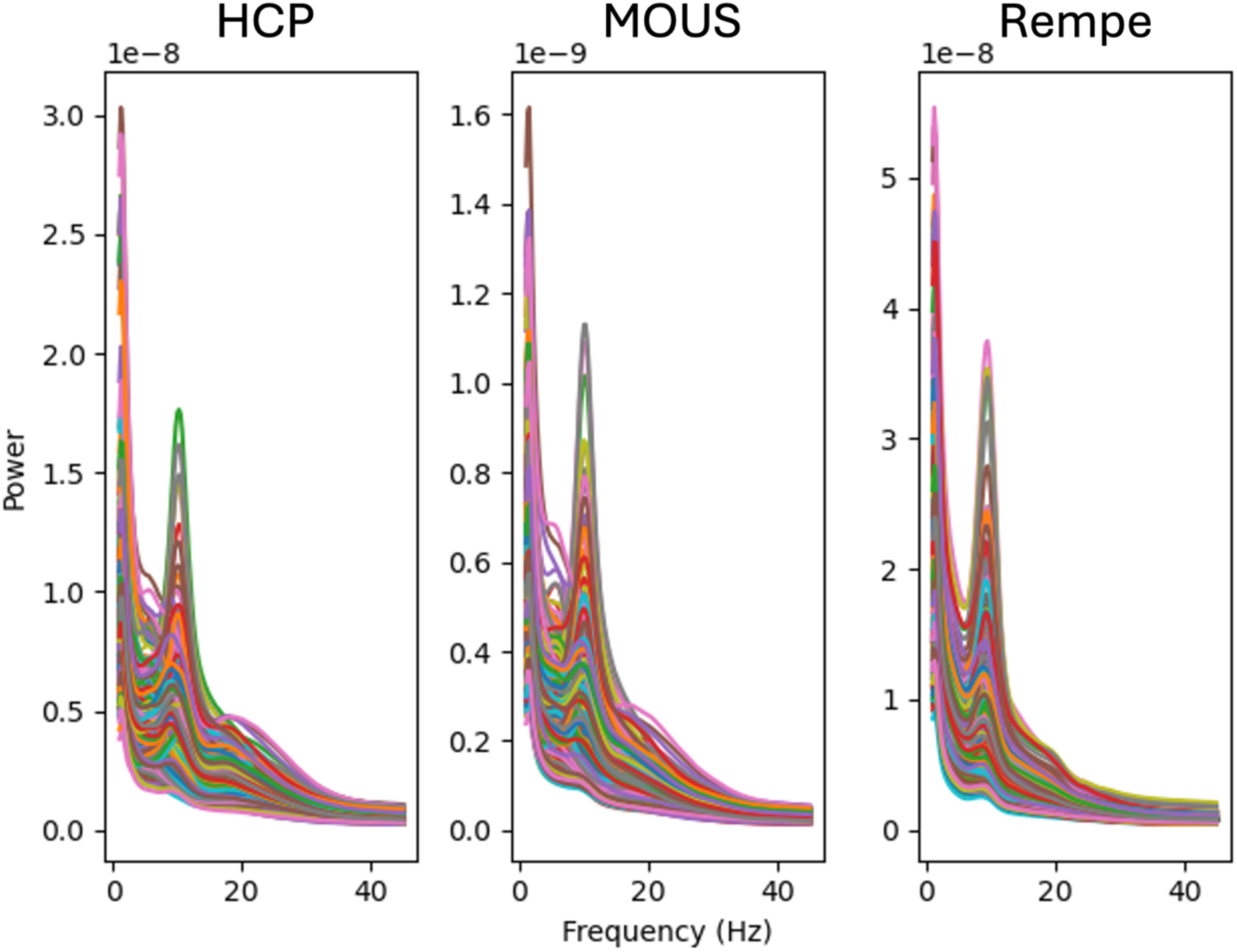
Power spectra averaged over all subjects for each of the three cohorts. Each line in the graph represents a single cortical parcel.

**Figure 6:**
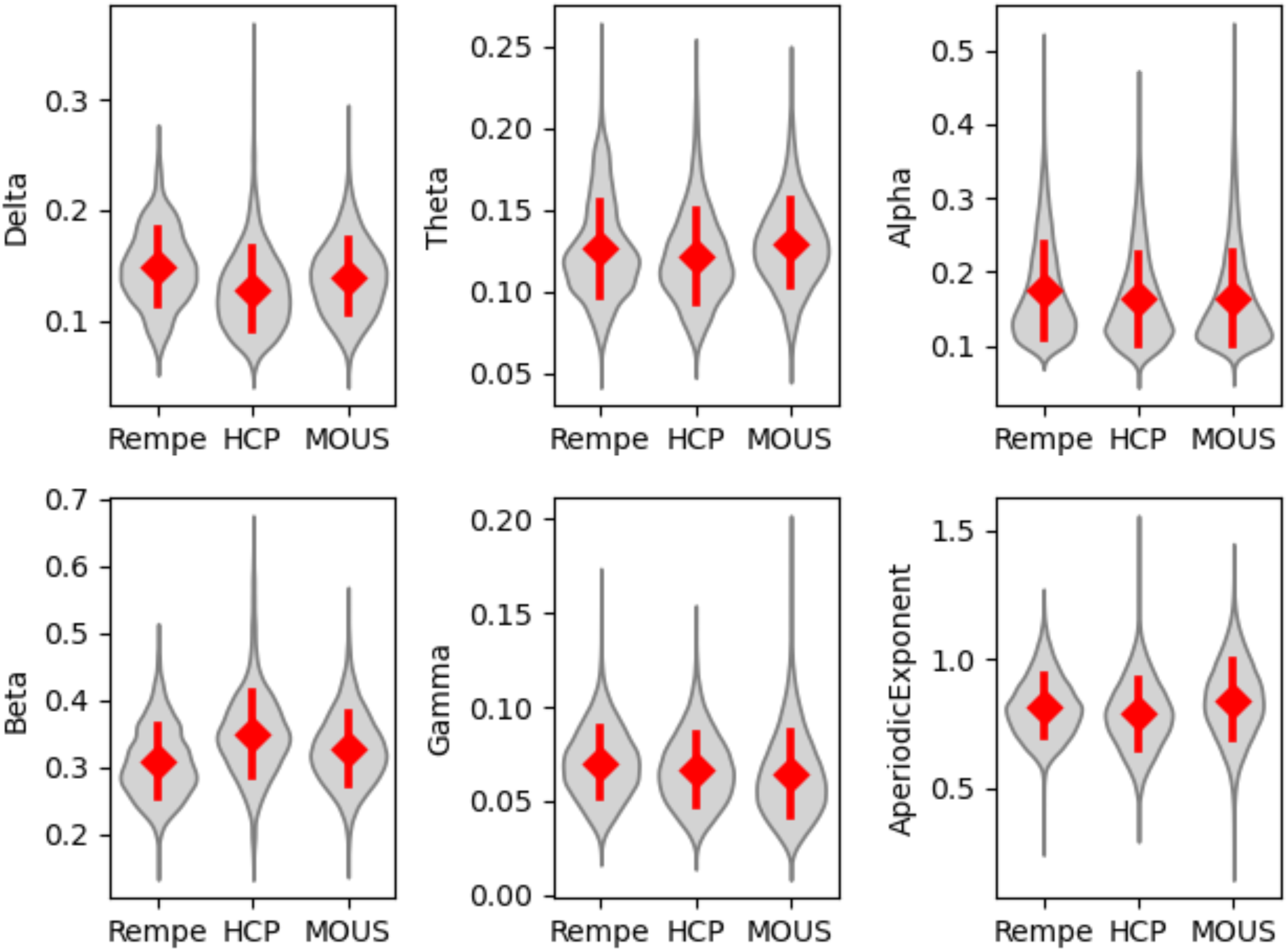
Distribution of data across parcels and subjects within each cohort. A subset of the Rempe cohort (aged 18-35) was used to better match the other two cohorts.

**Figure 7:**
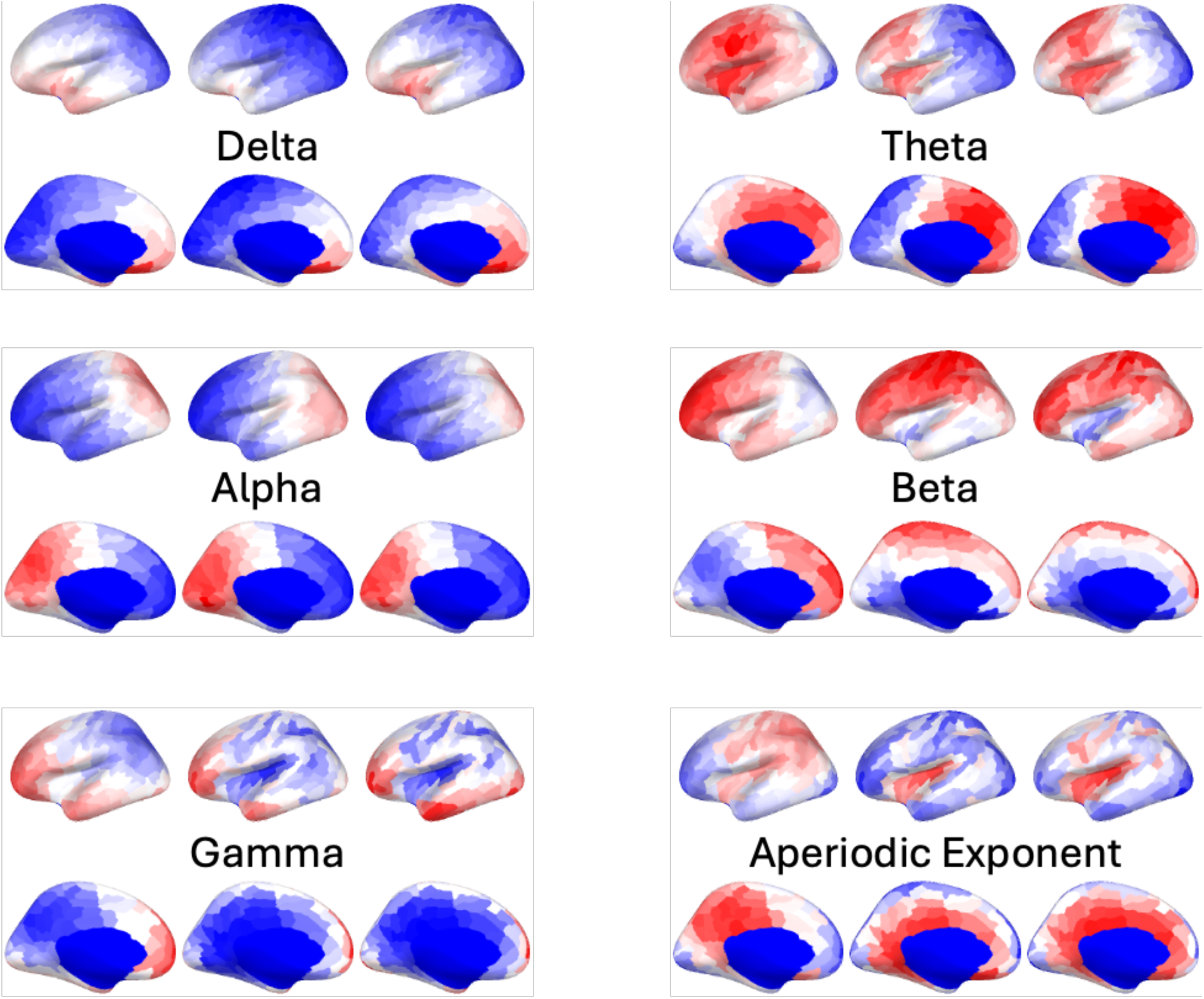
Plots of relative alpha power in the three datasets (with the aged 18-35 Rempe cohort), illustrating the familiar posterior to anterior gradient. All three datasets show remarkable similarity.

To obtain more qualitative values for overlap, we calculate Pearson correlation coefficients across all parcels between cohorts again using the truncated young-adult Rempe cohort, as a measure of spatial overlap. Correlation coefficients are given in Table 4. Finally, the HCP cohort included 81 participants that had both the first resting state run and a valid second resting state run. We calculate and report Intraclass correlation coefficients (ICC) for the spectral measures. In addition, to show spatial overlap, we also correlate the HCP cohort run 1 and run 2 power maps and report those values in Table 5.

**Table 4:** Correlations between spatial maps for each cohort, and for each spectral band, along with aperiodic exponent. Note that for the purposes of comparison, we selected a subset of the Rempe dataset in the same age range as the MOUS and HCP datasets.

|  | Spatial Correlations (all p-values < 5e-30) |  |  |
| --- | --- | --- | --- |
|  | Rempe vs HCP | Rempe vs MOUS | HCP vs MOUS |
| Delta | 0.925 | 0.923 | 0.970 |
| Theta | 0.774 | 0.867 | 0.928 |
| Alpha | 0.932 | 0.960 | 0.983 |
| Beta | 0.503 | 0.631 | 0.877 |
| Gamma | 0.779 | 0.789 | 0.919 |
| Aperiodic Exponent | 0.543 | 0.676 | 0.916 |

**Table 5:** Correlations between spatial maps for the first and second resting state runs for the HCP cohort. In addition, Intraclass correlation coefficients are given.

|  | Spatial Correlations<br>HCP Run1 vs Run2 | ICC(3,2)<br>HCP Run1 vs Run2 |
| --- | --- | --- |
| Delta | 0.995 | 0.759 |
| Theta | 0.966 | 0.847 |
| Alpha | 0.998 | 0.941 |
| Beta | 0.969 | 0.939 |
| Gamma | 0.998 | 0.932 |
| Aperiodic Exponent | 0.995 | 0.823 |

## Discussion

In this manuscript, we present a software pipeline designed as part of the ENIGMA MEG working group to enable automated, cortically localized, spectral analysis of large, multi-site and multi-platform datasets. Although the codebase is currently relatively inflexible (it is designed to produce power spectral density characteristics in brain parcels), it is modular in nature and thus easily extensible to other scientific aims and will be extended as ENIGMA MEG working group projects evolve. Notably, the key to automating our pipeline was a re-training of the MEGnet algorithm [34], so that accurate detection of artifacts could be accomplished without time consuming (and potentially biased) manual intervention. Finally, the use of a graphical QA interface enables high throughput review of data quality, vital to ensure the integrity of results.

Note that no harmonization of the cohorts we presented was performed in this demonstration, given that the goal was to show functionality rather than to test a specific hypothesis. Nevertheless, we would like to emphasize that harmonization is a crucial step for analysis of any multi-site data collection. To this end, code for harmonization of composite output files using a GAM extension of the ComBat algorithm (implemented in the NeuroHarmonize package [59]) is available in the enigmeg/group/ folder of the ENIGMA MEG pipeline code. Harmonization is performed on the final cohort after all data processing has been completed. Recently, we published a full exploration of harmonization for data processed using the ENIGMA MEG pipeline [60], which includes a comprehensive statistical evaluation of the effects of study cohort. We report F values for an ANOVA on the effect of study, as well as F values for Levene’s test of equality of variances across studies, and demonstrate that post-harmonization, the effect of study is not detectable. Given that the null hypothesis test framework is affected by study power and does not quantitatively show residual site variance, we also report partial R^2^ values attributable to the study effect and absolute log variance ratio, also demonstrating the efficacy of harmonization. Before harmonization, spectral maps in the five canonical bands showed median Pearson R correlations across 7 studies ranging from 0.80 to 0.94, while post-harmonization the range was 0.96-0.99. Median concordance correlation coefficients ranged from 0.59 to 0.89 before harmonization, and from 0.81 to 0.96 post-harmonization [60].

It is notable that even without data harmonization, there is substantial overlap in the distributions of the spectral power metrics across the three cohorts, both in the unaveraged data distributions across all subjects and all parcels, and in the spatial maps averaged over all subjects within a cohort. Note that these studies have different age distributions within the selected age range, and the Rempe dataset was acquired in an eyes closed condition while the other two cohorts were collected in an eyes open condition. Additionally, when comparing the test/re-test results for the HCP cohort, where data harmonization is not required, correlation values of spatial maps of spectral characteristics were extremely high (R > 0.96) for all metrics, and ICC values were good to excellent for all metrics.

The primary limitation of the ENIGMA pipeline is its relative inflexibility. However, the primary reason for the development of the ENIGMA pipeline was to enable source localized spectral analysis of multi-site resting state data in the context of the ENIGMA MEG working group. We demonstrate the construction of a pipeline that meets these needs, but is also modular and readily extensible to other analytic tasks. Referring to Figure 1, additional methods for alternative artifact cleaning methods could be inserted to provide cleaning of the data beyond MEGnet. While we chose the FOOOF/Specparam algorithm due to its simplicity and widespread usage, the spectral parameterization module could be replaced or augmented using an algorithm such as IRASA, which has a python implementation and can provide time resolved spectral estimates [61, 62], which could address additional scientific questions beyond the capabilities of the current pipeline. We anticipate that future ENIGMA MEG Working Group projects will continue to expand this platform, extending its use beyond spectral analysis of resting state MEG, and increasing its applicability to other resting-state related analyses (such as connectivity) and task-based MEG. While extension of the pipeline to task based data is possible, we would not expect that the degree of flexibility offered by large MEG analysis packages will be feasible in our framework. The addition of task-based analysis would likely be driven by a particular hypothesis and task, which we believe is an appropriate approach given the difficulty of assembling a very large dataset using a single MEG task. It would likely involve additional methods such as a “do event” event processing method, a separate “do_event_epochs” method specific to task based data, and a modified “do_task_beamformer” optimized for the particular task and hypothesis. Custom output methods would also be created. Using the existing ENIGMA Pipeline framework, however, would have the advantage of using the logging features, automated artifact correction, and high-throughput QA functionality present in the current pipeline.

Another potential limitation of the pipeline is the computational intensity. In particular, the reliance on FreeSurfer may impose a constraint for users with older hardware or without access to a computing cluster. There are alternatives to FreeSurfer, with the most notable being FastSurfer, a deep learning based tool that significantly speeds up the processing of datasets [63]. FastSurfer produces output files that are fully compatible with FreeSurfer, and could potentially be used as a drop-in alternative for users, although installation fo FreeSurfer will still be required for the ENIGMA MEG pipeline, as MNE python uses FreeSurfer functions to generate the bem surface. Recently, a number of processing platforms have appeared which use communal cloud computing resources to allow users to analyze large datasets online, including CBRAIN [64], brainlife.io [65], and the Dementias Platform UK (DPUK) [66]. These platforms reduce computational burden, but provide a restricted suite of analysis tools; however, the ENIGMA pipeline could be installed on these platforms for users without sufficient computational resources.

A final important issue to mention when combining datasets is coregistration accuracy. In the ENIGMA MEG pipeline, we chose to require the location of the MEG fiducial coils in the space of the anatomical MRI in the .json sidecar files as a prerequisite. If a head shape digitization is present, these fiducial points can be derived even if they are not in the shared data, although it is clearly preferable to have the creator of the dataset share the coregistration data they derived and used for their study. Although we provide QA images for assessing coregistration, it can be difficult to precisely judge coregistration accuracy when the face of the MRI is missing, as is standard practice for shared repositories. There additionally may be differences in coregistration accuracy across vendors or sites, with greater accuracy at sites employing head shape digitization.

Importantly, the development of the ENIGMA pipeline illustrates how software can be designed in keeping with FAIR principles. FAIR principles were developed to guide the sharing of scientific data in a way that is Findable, Accessible, Interoperable, and Reusable. Recently, analogous FAIR principles have been identified for research software [67] and are as follows. Findable software is registered in a permanent catalog or database with a globally unique and persistent identifier, such as a DOI. Findable software can also be easily found through a search and is indexed with metadata. Accessible software can be easily downloaded, with metadata available even when the software is no longer available. Interoperable software reads and writes data that is easily transferred between programs and conforms to domain-specific standards. Reusable software is licensed, clearly references other software packages, has a detailed revision history, and is well described. The ENIGMA pipeline has a DOI from Zenodo and is findable by internet search [35]. It can be easily downloaded from a GitHub repository, which also describes relevant metadata. The ENIGMA pipeline requires data in BIDS format for input, which is the neuroimaging community standard for data sharing. Output data utilizes the BIDS derivative structure, and industry standard file formats or simple .csv files. The software is in the public domain, which is clearly described in the “License” section of the GitHub repository. The code includes dependencies to other Python libraries and algorithms as referenced on the GitHub page and herein.

The MEG research community would benefit from more collaborative, multi-site studies to enhance reliability, generalizability, and statistical power. To carry out studies of this nature, however, tools specifically designed for large scale data analysis are needed. While the ENIGMA MEG pipeline does not currently meet all data analysis needs, it paves the way for further development of automated, high-throughput, software pipelines developed with FAIR principles in mind.

## Data Availability

The following datasets were used:

Human Connectome Project data is available through ConnectomeDB after agreeing to Open Access Data Use Terms: https://db.humanconnectome.org/.

MOUS data is available through the Radboud University Data Repository after signing a Data Use Agreement: https://doi.org/10.34973/37n0-yc51.

The Rempe dataset is open access and can be downloaded through the Boys Town National Research Hospital: https://cdn.boystown.org/media/Rempe_Ott_PNAS_2023_Data.zip.

## Source Code Availability and Requirements

Project Name: ENIGMA MEG pipeline

Project homepage: https://github.com/nih-megcore/enigma_MEG

Software documentation: https://github.com/nih-megcore/enigma_MEG

Operating system(s): Linux

Programming language: Python

Other Requirements: Python (<3.12), FreeSurfer, MNE Python (1.5)

License: Unlicense (public domain)

## Authors’ contributions

ACN: Conceptualization, Data curation, Formal analysis, Methodology, Resources, Software, Supervision, Validation, and Writing – original draft.

AMN: Data curation, Project administration, Validation, and Writing – review and editing.

FWC: Conceptualization, Methodology, and Writing – review and editing.

PMT: Conceptualization and Writing – review and editing.

JDS: Conceptualization, Data curation, Formal Analysis, Methodology, Software, Validation, and Writing – review and editing.

## Acknowledgements

The NIH ENIGMA MEG working group team appreciates the continued support of the USC ENIGMA team. This work utilized the computational resources of the NIH HPC Biowulf Cluster.

