## Supplemental Materials for "The ENIGMA MEG Pipeline: Automated cortically localized spectral analysis of multi-site resting state MEG datasets"

Supplementary Tables 1-4: Confusion matrices from MEGnet retraining grouped by site/MEG vendor

| | Predicted Class - HCP | | | | | | |
|---|---|---|---|---|---|---|---|
| **Actual Class** | | Non-Artifact | Blink | Cardiac | Saccade | **Sensitivity** | **False Negative Rate** |
| | Non-Artifact | 517 | 2 | 9 | 0 | 97.92% | 2.08% |
| | Blink | 0 | 32 | 0 | 0 | 100.00% | 0.00% |
| | Cardiac | 0 | 0 | 39 | 0 | 100.00% | 0.00% |
| | Saccade | 0 | 0 | 0 | 21 | 100.00% | 0.00% |
| | **Positive Predictive Value** | 100.00% | 94.12% | 81.25% | 100.00% | **Accuracy** | 98.23% |
| | **False Discovery Rate** | 0.00% | 5.88% | 18.75% | 0.00% | **Error Rate** | 1.77% |

| | Predicted Class - NYU | | | | | | |
|---|---|---|---|---|---|---|---|
| **Actual Class** | | Non-Artifact | Blink | Cardiac | Saccade | **Sensitivity** | **False Negative Rate** |
| | Non-Artifact | 168 | 0 | 3 | 1 | 97.67% | 2.33% |
| | Blink | 0 | 10 | 0 | 0 | 100.00% | 0.00% |
| | Cardiac | 0 | 0 | 14 | 0 | 100.00% | 0.00% |
| | Saccade | 0 | 0 | 0 | 4 | 100.00% | 0.00% |
| | **Positive Predictive Value** | 100.00% | 100.00% | 82.35% | 80.00% | **Accuracy** | 98.00% |
| | **False Discovery Rate** | 0.00% | 0.00% | 17.65% | 20.00% | **Error Rate** | 2.00% |

| | Predicted Class - CAMCAN | | | | | | |
|---|---|---|---|---|---|---|---|
| **Actual Class** | | Non-Artifact | Blink | Cardiac | Saccade | **Sensitivity** | **False Negative Rate** |
| | Non-Artifact | 1040 | 2 | 7 | 2 | 98.95% | 1.05% |
| | Blink | 0 | 58 | 0 | 0 | 100.00% | 0.00% |
| | Cardiac | 0 | 0 | 56 | 0 | 100.00% | 0.00% |
| | Saccade | 0 | 0 | 0 | 15 | 100.00% | 0.00% |
| | **Positive Predictive Value** | 100.00% | 96.67% | 88.89% | 88.24% | **Accuracy** | 99.07% |
| | **False Discovery Rate** | 0.00% | 3.33% | 11.11% | 11.76% | **Error Rate** | 0.93% |

| | Predicted Class - NIH | | | | | | |
|---|---|---|---|---|---|---|---|
| **Actual Class** | | Non-Artifact | Blink | Cardiac | Saccade | **Sensitivity** | **False Negative Rate** |
| | Non-Artifact | 1460 | 5 | 9 | 5 | 98.72% | 1.28% |
| | Blink | 0 | 79 | 0 | 0 | 100.00% | 0.00% |
| | Cardiac | 0 | 0 | 86 | 0 | 100.00% | 0.00% |
| | Saccade | 0 | 0 | 0 | 36 | 100.00% | 0.00% |
| | **Positive Predictive Value** | 100.00% | 94.05% | 90.53% | 87.80% | **Accuracy** | 98.87% |
| | **False Discovery Rate** | 0.00% | 5.95% | 9.47% | 12.20% | **Error Rate** | 1.13% |

Supplementary Tables 5-7: Confusion matrices from MEGnet retraining grouped by eyes open rest, eyes closed rest, and task

| | Predicted Class - Rest Eyes Open | | | | | | |
|---|---|---|---|---|---|---|---|
| Actual Class | | Non-Artifact | Blink | Cardiac | Saccade | **Sensitivity** | **False Negative Rate** |
| | Non-Artifact | 393 | 0 | 6 | 1 | 98.25% | 1.75% |
| | Blink | 0 | 23 | 0 | 0 | 100.00% | 0.00% |
| | Cardiac | 0 | 0 | 30 | 0 | 100.00% | 0.00% |
| | Saccade | 0 | 0 | 0 | 7 | 100.00% | 0.00% |
| | **Positive Predictive Value** | 100.00% | 100.00% | 83.33% | 87.50% | **Accuracy** | 98.48% |
| | **False Discovery Rate** | 0.00% | 0.00% | 16.67% | 12.50% | **Error Rate** | 1.52% |

| | Predicted Class - Rest Eyes Closed | | | | | | |
|---|---|---|---|---|---|---|---|
| Actual Class | | Non-Artifact | Blink | Cardiac | Saccade | **Sensitivity** | **False Negative Rate** |
| | Non-Artifact | 386 | 0 | 2 | 0 | 99.48% | 0.52% |
| | Blink | 0 | 19 | 0 | 0 | 100.00% | 0.00% |
| | Cardiac | 0 | 0 | 24 | 0 | 100.00% | 0.00% |
| | Saccade | 0 | 0 | 0 | 9 | 100.00% | 0.00% |
| | **Positive Predictive Value** | 100.00% | 100.00% | 92.31% | 100.00% | **Accuracy** | 99.55% |
| | **False Discovery Rate** | 0.00% | 0.00% | 7.69% | 0.00% | **Error Rate** | 0.45% |

| | Predicted Class -Task | | | | | | |
|---|---|---|---|---|---|---|---|
| Actual Class | | Non-Artifact | Blink | Cardiac | Saccade | **Sensitivity** | **False Negative Rate** |
| | Non-Artifact | 2406 | 9 | 20 | 7 | 98.53% | 1.47% |
| | Blink | 0 | 137 | 0 | 0 | 100.00% | 0.00% |
| | Cardiac | 0 | 0 | 141 | 0 | 100.00% | 0.00% |
| | Saccade | 0 | 0 | 0 | 60 | 100.00% | 0.00% |
| | **Positive Predictive Value** | 100.00% | 93.84% | 87.58% | 89.55% | **Accuracy** | 98.71% |
| | **False Discovery Rate** | 0.00% | 6.16% | 12.42% | 10.45% | **Error Rate** | 1.29% |

Supplementary Figure 1: Distribution of ages for the three cohort datasets.

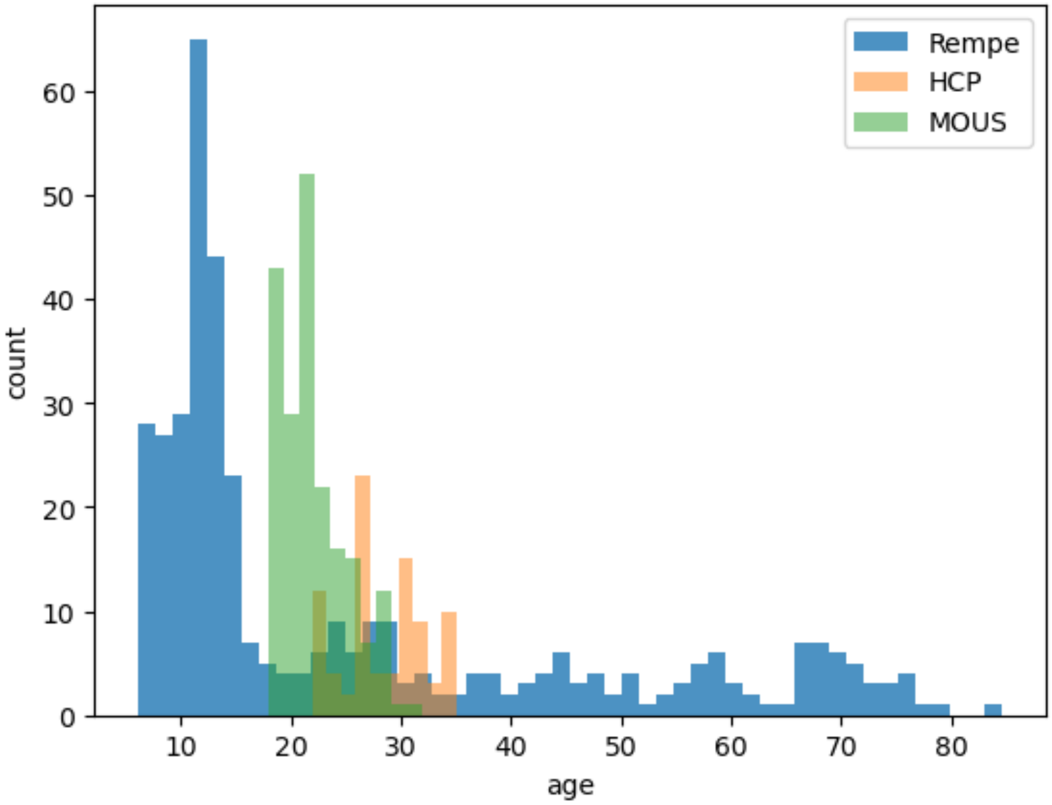
